# Systematic Mendelian randomization framework elucidates hundreds of genetic loci which may influence disease through changes in DNA methylation levels

**DOI:** 10.1101/189076

**Authors:** Tom G. Richardson, Philip C. Haycock, Jie Zheng, Nicholas J. Timpson, Tom R. Gaunt, George Davey Smith, Caroline L. Relton, Gibran Hemani

**Affiliations:** MRC Integrative Epidemiology Unit (IEU), Bristol Medical School (Population Health Sciences), University of Bristol, Oakfield House, Oakfield Grove, Bristol, BS8 2BN, United Kingdom

## Abstract

We have undertaken an extensive Mendelian randomization (MR) study using methylation quantitative trait loci (mQTL) as genetic instruments to assess the potential causal relationship between genetic variation, DNA methylation and 139 complex traits. Using two-sample MR, we observed 1,191 effects across 62 traits where genetic variants were associated with both proximal DNA methylation (i.e. cis-mQTL) and complex trait variation (P<1.39x10^−08^). Joint likelihood mapping provided evidence that the causal mQTL for 364 of these effects across 58 traits was also likely the causal variant for trait variation. These effects showed a high rate of replication in the UK Biobank dataset for 14 selected traits, as 121 of the attempted 129 effects replicated. Integrating expression quantitative trait loci (eQTL) data suggested that genetic variants responsible for 319 of the 364 mQTL effects also influence gene expression, which indicates a coordinated system of effects that are consistent with causality. CpG sites were enriched for histone mark peaks in tissue types relevant to their associated trait and implicated genes were enriched across relevant biological pathways. Though we are unable to distinguish mediation from horizontal pleiotropy in these analyses, our findings should prove valuable in identifying candidate loci for further evaluation and help develop mechanistic insight into the aetiology of complex disease.

## Background

The majority of genetic variants associated with complex traits are located in non-coding regions of the genome and therefore likely to influence disease via gene regulation(Edwards et al., 2013). To develop our understanding of these mechanisms, studies have incorporated data concerning genetic variants associated with gene expression into analyses (also known as expression quantitative trait loci (eQTL)(Zhu et al., 2016, Burkhardt et al., 2015, Mancuso et al., 2017). Recently, this type of methodology has been extended to integrate epigenetic data using genetic variants associated with DNA methylation levels (known as methylation quantitative trait loci (mQTL)) (Hannon et al., 2017, Richardson et al., 2017). In this study, we have built on this previous work to comprehensively investigate whether DNA methylation plays a mediatory role along the causal pathway from genetic variation to complex trait and disease susceptibility.

As with complex traits, DNA methylation levels at CpG sites across the genome can be determined by both genetic and environmental factors. Moreover, both complex traits and DNA methylation are prone to confounding and reverse causation, which can undermine our ability to infer causal relationships (McRae et al., 2014, Relton and Davey Smith, 2010). An approach to address this limitation is Mendelian randomization (MR), a method by which the causal inference of one trait (the exposure) on another trait (the outcome) can be inferred. This is achieved by using genetic variants known to robustly associate with the exposure as instrumental variables (Davey Smith and Hemani, 2014, Davey Smith and Ebrahim, 2003). The sample size of studies with data on epigenome-wide DNA methylation, genome-wide genetic data and complex traits are modest compared to most genetic association studies of complex traits, primarily due to the current costs of DNA methylation arrays. A recent methodological development to circumvent this limitation is two-sample MR (2SMR), an approach where summary statistics for the observed effect of genetic instruments on exposure and outcome are obtained from two separate studies (Burgess et al., 2015, Pierce and Burgess, 2013). In doing so, causal relationships can be investigated without requiring a sample of individuals with genotype, exposure and outcome data.

As described in our previous work (Richardson et al., 2017), when a genetic variant is reliably associated with both DNA methylation and complex trait variation, we postulate that there are 4 possible scenarios that may account for this (Figure 1):

**Figure 1:**
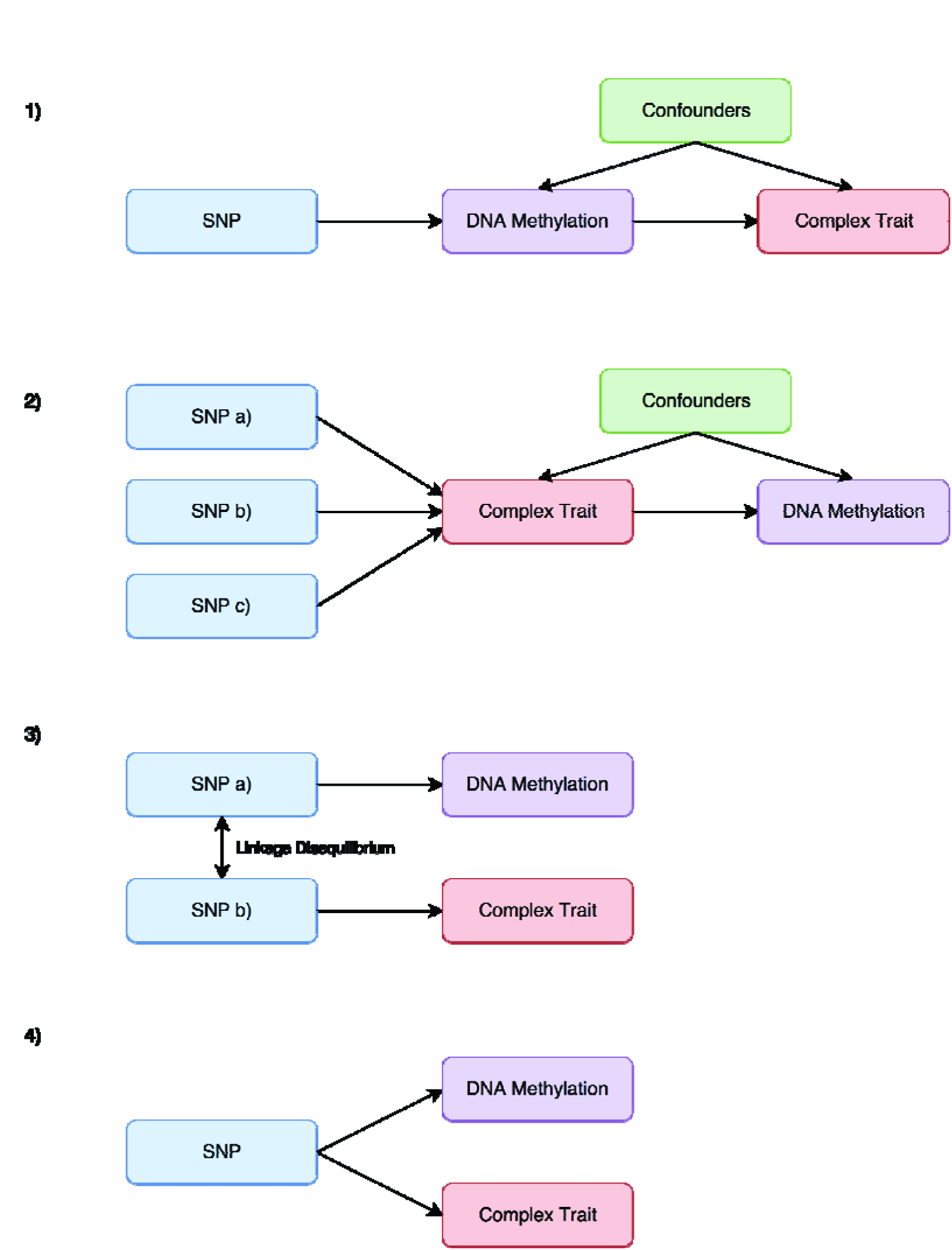
Explanations evaluated which may potentially explain observed associations between methylation quantitative trait loci and trait outcomes. 1) The genetic variant has a causal effect on the complex trait which is mediated by changes in DNA methylation. 2) The genetic variant has a causal effect on the complex trait which subsequently influences DNA methylation at this locus. 3) The genetic variant that influences DNA methylation is in linkage disequilibrium (LD) with another variant that influences complex trait variation. 4) The genetic variant influences DNA methylation and the complex trait via two independent biological pathways (also known as horizontal pleiotropy).

1. The genetic variant has a causal effect on the complex trait which is mediated by changes in DNA methylation.
2. The genetic variant has a causal effect on the complex trait (or a related complex trait which resides along the causal pathway to disease) which subsequently influences DNA methylation at this locus.
3. The genetic variant responsible for changes in DNA methylation is in linkage disequilibrium (LD) with the genetic variant that influences complex trait variation.
4. The genetic variant influences DNA methylation and the complex trait via two independent biological pathways (also known as horizontal pleiotropy).

Within our analytical framework, we first attempt to distinguish between explanations 1 and 2 by using 2SMR to evaluate the causal influence of DNA methylation on complex traits and then conversely the opposite direction of effect (also known as bi-directional MR (Timpson et al., 2011, Vimaleswaran et al., 2013)). A limitation of this approach is that DNA methylation can only typically be instrumented by a single cis-acting variant, which means that an unreliable MR estimate of causality may arise due to the causal variant for DNA methylation simply being in linkage disequilibrium with the causal trait variant (explanation 3). The chances of this occurrence is dramatically increased when investigating causal relationship systematically as undertaken in our framework. A potential approach to mitigate this limitation is using a colocalization approach, such as the joint likelihood mapping (JLIM) method. This approach has been devised to investigate whether the underlying genetic variation at a genomic region is responsible for observed effects on both an intermediate and complex trait (Chun et al., 2017).

A single cis-acting instrument also means that we are unable to reliably distinguish between mediation (explanation 1) and horizontal pleiotropy (explanation 4). Nevertheless, within our framework we use MR to investigate the relationship between DNA methylation and gene expression at loci where mediation is a potential explanation of observed effects. In doing so, we aim to identify a coordinated system of effects that are consistent with causality, such as genetic variants influencing gene expression via changes in DNA methylation.

In this study, we have adapted our analytical framework developed previously to evaluate the causal relationship between DNA methylation and 139 complex traits taken from large-scale consortia using a two-sample framework (Hemani et al., 2016). We build on previous work (Hannon et al., 2017) by extending the survey to a much larger number of traits, interrogating bi-directional relationships, integrating gene expression data into analyses and undertaking exhaustive joint likelihood mapping analyses to investigate linkage as an explanation for observed effects. Validation of results with evidence of a causal relationship for a selection of traits was undertaken using data from up to 334,398 individuals enrolled in the UK Biobank study (Sudlow et al., 2015). Functional annotation and enrichment analyses, including data for histone mark peaks and DNAse I hypersensitivity sites across 113 different tissue types, was undertaken for selected variants and CpG sites (Romanoski et al., 2015, Encode Project Consortium et al., 2007).

## Results

### Systematic evaluation of the causal relationship between DNA methylation and complex traits

The initial analysis involved over 4.2 million MR analyses to evaluate the causal relationship between DNA methylation at 30,328 CpG sites and 139 complex traits using MR-Base. A list of these traits can be found in Supplementary Table 1, which were selected based on the sample size and population analysed in their respective GWAS. We only investigated CpG sites using cis-mQTL (i.e. genetic instruments within 1MB distance of their associated CpG site) in order to reduce the risk of pleiotropy influencing our results. Subsequently the majority of CpG sites were instrumented using a single cis-acting mQTL (n=26,975) and therefore MR effect estimates were calculated using the Wald ratio. When more than one instrument was available the inverse variance weighted (IVW) method was used instead.

There were 1,191 observed effects (i.e. associations between a CpG site and complex trait) which survived the multiple testing threshold across 62 different traits (P < 1.397 × 10^−08^, Supplementary Table 2). This threshold was based on the number of tests undertaken across independent traits using the PhenoSpD method (Zheng et al., 2017, Nyholt, 2004, Cichonska et al., 2016)). CpG sites were annotated based on evaluations of the Illumina 450K array (Naeem et al., 2014, Zhou et al., 2017). A heat map visualising the correlation of the z scores from the MR analysis across traits can be found in Supplementary Fig. 1, which highlights traits which may be influenced by changes in DNA methylation at shared loci. Figure 2 provides an overview of the analysis pipeline applied in this study for downstream analyses concerning these results.

**Figure 2:**
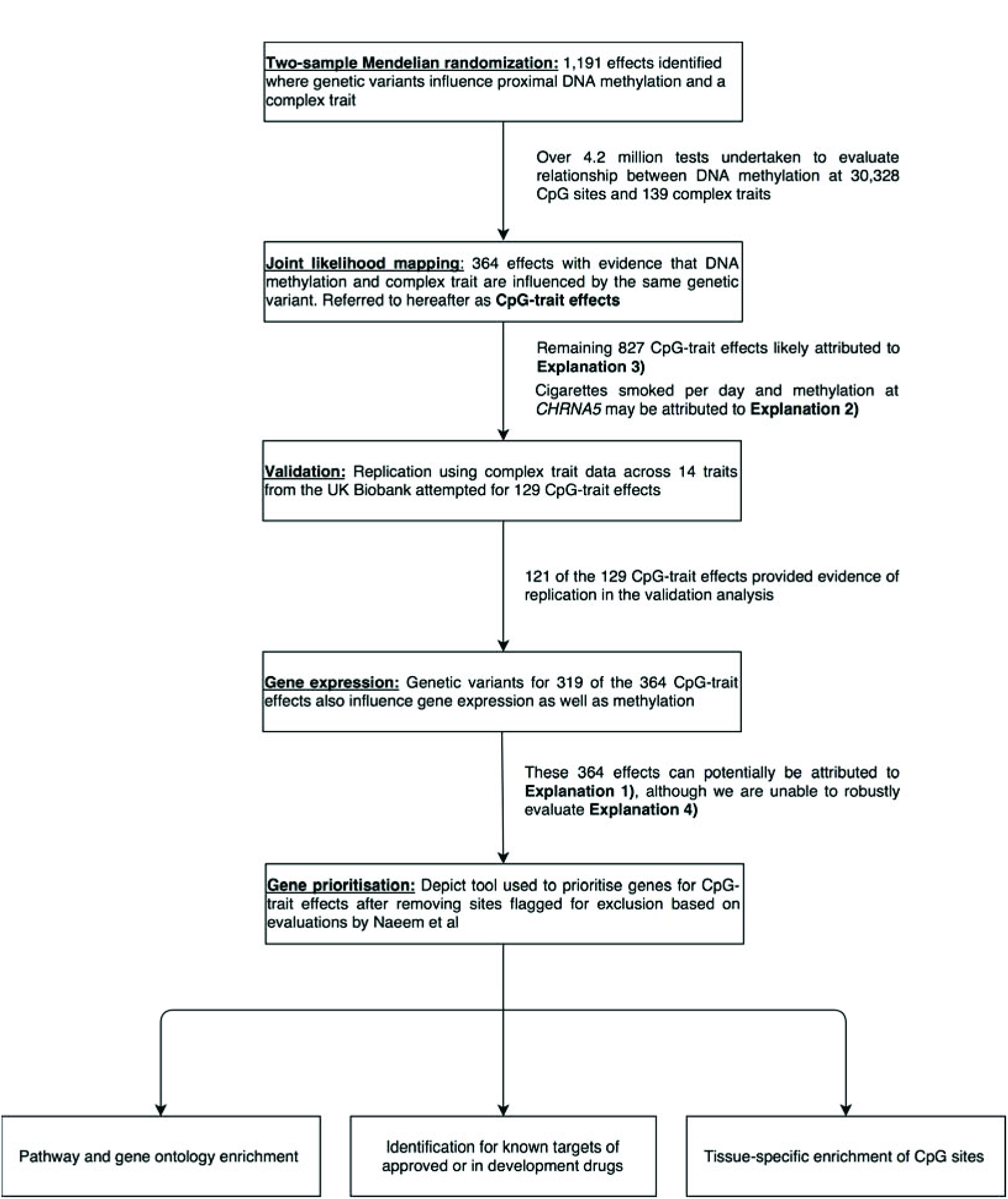
Analysis pipeline to evaluate explanations for observed associations between methylation quantitative trait loci and trait outcomes. This flowchart provides an overview of the analysis plan in this study to evaluate 4 different explanations which may explain trait-associated methylation quantitative trait loci. Explanations 1 to 4 are as described in Figure 1.

### Identifying causal variants for both DNA methylation and complex traits

Results surviving multiple testing in the previous analysis may arise due to an mQTL and trait-associated variant overlapping at a genomic locus due to chance. To investigate this, we applied the JLIM algorithm (Chun et al., 2017) which tests whether variation in two traits (i.e. DNA methylation and a complex trait in this study) are driven by a shared causal effect. This is ascertained by generating a permutation-based null distribution for a trait with individual-level data (i.e. DNA methylation in our analysis) and assessing the likelihood that the causal variant for this trait is also responsible for variation on a different trait based on summary-level data (i.e. GWAS results for a complex trait). Permutation testing was implemented by the JLIM method to account for the 1,191 effects identified in the previous analysis (P < 4.20 × 10^−5^). The JLIM results suggested that 364 of the 1,191 CpG-trait effects were observed due to methylation and complex trait variation both being influenced by the same underlying genetic variant (Supplementary Table 3). We refer to these 364 effects hereafter as ‘CpG-trait effects’ as they represent associations where DNA methylation may reside along the causal pathway from genetic variant to complex trait.

Consequently, the 805 effects which did not provide evidence from joint likelihood mapping in this evaluation were likely observed due to the causal variant for DNA methylation being in linkage disequilibrium with a separate variant responsible for complex trait variation. Figure 3 illustrates findings for 2 of the 62 traits which had at least one effect that survived the multiple testing threshold, where individual points represent p-values from the 2SMR analysis. Interpretation of these findings are different to those illustrated by a conventional Manhattan plot in a GWAS. For instance, using the strongest observed effect in Figure 3 as an example, a standard deviation increase in DNA methylation at the *SLC12A4* locus results in a 0.138 standard deviation decrease in HDL cholesterol (and vice versa). Points highlighted in red correspond to loci where the JLIM provided evidence that the same underlying causal variant influences both DNA methylation and complex trait. Manhattan plots for all 62 traits can be found in Supplementary File 1.

**Figure 3:**
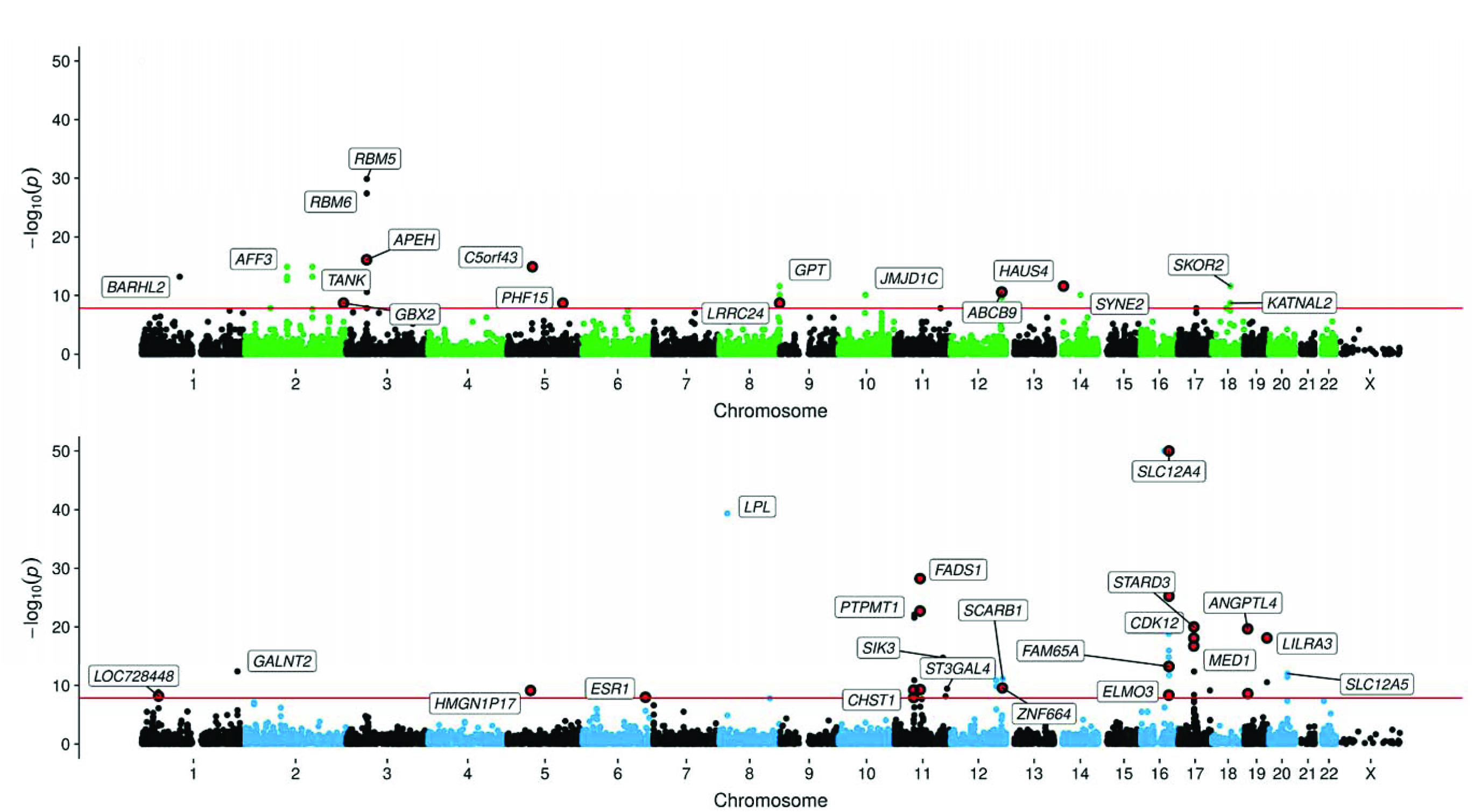
Manhattan plots illustrating results of two-sample Mendelian randomization analysis between epigenome-wide DNA methylation and a) educational attainment (top) b) high density lipoprotein cholesterol (bottom). Points represent −log10 p-values (y-axis) for CpG sites (genomic location on the x-axis) as evaluated using two-sample Mendelian randomization analysis between DNA methylation (as our exposure) and complex traits (as our outcome) using mQTL as genetic instruments. Effects that survive the multiple testing threshold in our analysis (P<1.397 × 10^−08^ – represented by the red horizontal line) are annotated using mapped genes according to Illumina (or nearest gene when no gene has been reported by Illumina). Effects where joint likelihood mapping suggested the causal variant for DNA methylation and complex trait variation were the same are highlighted in red.

### Reverse Mendelian randomization

For the 364 CpG-trait effects identified in the previous analysis, we undertook reverse MR to evaluate evidence of genetic liability between complex traits and DNA methylation. This was undertaken by modelling a complex trait as our exposure and DNA methylation levels at a CpG as our outcome. The only evidence of liability was observed between number of cigarettes smoked per day and DNA methylation variation at the *CHRNA5/PSMA4* region (Supplementary Table 4). However, this complex trait currently only has a single genetic instrument which weakens our ability to robustly investigate direction of effect for this result.

More broadly, all results from the reverse MR analysis should be interpreted with caution, as we do not have data on complex trait incidence. Therefore results can only be regarded as an association of the disease/trait liability as opposed to causality. For example, it is unlikely that incidence of coronary heart disease would have been frequent enough in the sample used to generate effect estimates on DNA methylation to identify a true causal effect. Furthermore, within a 2SMR framework, statistical power is determined by the sample size used to generate effect estimates on the outcome variable. Therefore, we may lack statistical power when modelling DNA methylation as our outcome variable, as samples sizes with DNA methylation were relatively modest compared to large-scale GWAS (n=~800). Nonetheless, this aspect of our framework is important to assess evidence of disease liability and should prove valuable as samples with DNA methylation data increases.

### Validation of findings within the UK Biobank

We undertook validation analyses for 129 of the CpG-trait effects using complex trait data from the UK Biobank (Supplementary Table 5) (Sudlow et al., 2015). There was evidence of validation for 121 of the 129 effects (P < 3.88 × 10^−04^, Supplementary Table 6), although all observed effects had P < 0.075 and also consistent directions of effect with DNA methylation as observed in the discovery analysis.

### Evaluating the relationship between DNA methylation and gene expression

We integrated gene expression data to investigate whether the genetic variants used to identify CpG-trait effects were known to influence gene expression as well as DNA methylation. Data from the GTEx consortium (Carithers and Moore, 2015) and the blood eQTL browser(Westra et al., 2013) suggested that this was the case for 319 of the 364 CpG-trait effects. 2SMR was used to evaluate the relationship between DNA methylation and gene expression at each of these loci i.e. whether an increase in DNA methylation results in either an increase or decrease in gene expression (Supplementary Table 7).

### Gene prioritisation, implicated biological pathways and druggable targets

A suite of bioinformatics tools was used to calculate the predicted consequences and severity for genetic variants responsible for CpG-trait effects (Supplementary Table 8). At this stage, any CpG sites recommended for exclusion based on evaluations of the 450K array(Naeem et al., 2014) (as annotated in Supplementary Table 2) were excluded from all further downstream analyses to remove any potential bias incurred by including them. Likely impacted genes for CpG-trait effects were determined using the gene prioritisation algorithm from DEPICT (Data-driven Expression-Prioritized Integration for Complex Traits) (Pers et al., 2015). When DEPICT was unable to identify a likely impacted gene we used the nearest gene instead (Supplementary Table 9). Annotated genes were then grouped into categories based on their associated trait (Supplementary Table 10). Each group of genes was then analysed in turn using ConsensusPathDB (Kamburov et al., 2013) to test whether likely implicated genes were enriched for biological pathways (Supplementary Table 11) and gene ontology terms (Supplementary Table 12) based on a false discovery rate < 5%. Overall there were 67 enriched pathway effects and 312 enriched GO term effects.

Prioritised genes were also evaluated for druggability using the ChEMBL database (Bento et al., 2014) (version 23 accessed on 13^th^ June 2017). Proteins encoded by implicated genes which are targets for therapeutic intervention were identified (Supplementary Table 13). These included approved drugs, such as estropipate and estradiol cypionate, which target *ESR1*, as well as compounds in development, such as cyclin-dependent kinase inhibitors, which target *CDK12*.

### Tissue specific enrichment for CpG sites

CpG sites implicated in CpG-trait effects were annotated to determine whether they reside in regulatory regions using data from Illumina and Ensembl (Yates et al., 2016). DNAse I and histone mark peak data across 113 different tissue types from the ENCODE and the Roadmap Epigenomics projects was also used to annotate CpG sites (Romanoski et al., 2015, Encode Project Consortium et al., 2007). CpG sites were then grouped according to the category of their associated trait (Supplementary Table 10) and tested for enrichment after removing proximal probes which may be co-methylated (Supplementary Tables 14-22). In particular, evidence of enrichment for H3K4me1 histone marks was observed for associated CpG sites, as well as evidence of enrichment in tissue types relevant for associated traits. For instance, the top hit for autoimmune traits was ob served for H3K4me1 marks in spleen tissue, whereas the top hit for haematological traits was observed for H3K4me1 marks in primary haematopoietic cells. Heat maps illustrating these results for histone mark peaks across different tissue types can be found in Supplementary Fig. 2a-g.

## Discussion

In this study we have extended an analytical framework to systematically evaluate the causal relationship between DNA methylation and complex traits using GWAS summary data. We identified 364 effects where genetic variants may be influencing disease via epigenetic processes. Although we are unable to robustly demonstrate that these effects occur along a common causal pathway to disease (e.g. the associations could be compatible with horizontal pleiotropy), we observed evidence that gene expression may also be influenced by genetic variants for 319 of these effects, suggesting a coordinated system that is consistent with causality. The genes impacted by changes in DNA methylation at these CpG sites represent promising candidates to explore the potential mediatory role of epigenetic modifications and their potential downstream effects on disease aetiology.

An attractive advantage of using 2SMR to investigate this relationship is that it circumvents the requirement of having both intermediate and complex traits measured in the same sample. For instance, a recent epigenome-wide association study (EWAS) of lipids used a sample size of 725 individuals in their discovery analysis to identify 2 CpG sites associated with HDL cholesterol. However, as illustrated in the bottom plot of Figure 3, using findings from a large-scale genetic association study (with approximately 190,000 individuals) we have discovered 9 genetic loci (which are different to the 2 identified in the aforementioned EWAS), which may influence HDL cholesterol variation via changes in DNA methylation. Furthermore, by using genetic instruments we are also less at risk of confounding and reverse causation biasing results. An example of this can be found by contrasting the top plot in Figure 3 with results from a recent EWAS of educational attainment, which identified associations at 9 CpG sites that were all previously associated with cigarette smoking (Linnér, et al. 2017). Although educational attainment may be an underlying cause of these changes in methylation levels (i.e. educational attainment influences smoking behaviour), such claims cannot be made with confidence in the presence of confounding factors. In contrast, none of the 7 independent CpG sites linked with educational attainment in this study are associated with exposure to cigarette smoking. This is based on findings from the largest smoking EWAS to date of both own smoking (Joehanes et al., 2016) and exposure to maternal smoking in utero (Joubert et al., 2016).

Integrating multiple types of ‘omic’ data into study designs is likely to become increasingly popular in the forthcoming years as the technologies required to generate data at scale become more feasible. Moreover, advancements in such technologies should allow a further detailed examination of the role of intermediate phenotypes in complex trait variation. For instance, the 450K Illumina Infinium Beadchip array used to generate the DNA methylation data in this study only covers ~1.7% of the 29 million CpG sites across the human genome (Ma et al., 2013). This suggests that a wealth of unmeasured data remains unexplored within this paradigm. Furthermore, although we have demonstrated the value of our analytical framework to investigate the role of DNA methylation in disease, we anticipate future studies will have success by investigating other intermediate traits in a similar manner, such as histone marks, metabolites and proteins. These endeavours will be valuable in uncovering signals which reflect a coordinated system of causality, as well as helping pinpoint the true causal gene at densely populated gene neighbourhoods. They should also prove particularly valuable to help identify and evaluate targets for therapeutic intervention.

Studies with increasingly large sample sizes with ‘omic’ data will also allow more robustly associated QTL across different omics types to be uncovered across the genome. This will be hugely beneficial for frameworks such as the one portrayed in this study as it should improve causal inference amongst intermediate traits and downstream implications on disease susceptibility. Moreover, using multiple instruments can improve our ability to separate mediation from horizontal pleiotropy as the putative mechanism underlying the association (Bowden et al., 2015, Bowden et al., 2016, Hartwig et al., 2017). The integration of co-localization methods to assess whether changes in DNA methylation and complex traits are driven by shared causal variants will remain important to implement. In this study, we have been able to use the JLIM method due to having individual level data on epigenome-wide DNA methylation from the ARIES project. Future endeavours, which may be restricted to using summary-level data for omics trait, are able to utilise viable alternatives, such as the HEIDI (heterogeneity in dependent instruments)(Zhu et al., 2016) and ‘coloc’ (Giambartolomei et al., 2014) methods.

The results presented in this study are likely only the tip of the iceberg for candidate loci which may influence complex traits via epigenetic mechanisms. Thorough evaluations of these loci are necessary to determine the extent to which these processes play a role in complex disease risk. A wealth of data on intermediate omic traits are expected to be generated in large sample sizes across multiple tissue types in the forthcoming years. Mendelian randomization can be used to interrogate causal relationships amongst these intermediate traits and help develop our understanding of the causal pathway from genetic variation to disease.

## Online methods

### The Avon Longitudinal Study of Parents and Children (ALSPAC)

ALSPAC is a population-based cohort study investigating genetic and environmental factors that affect the health and development of children. The study methods are described in detail elsewhere (Boyd et al., 2013, Fraser et al., 2013) (http://www.bristol.ac.uk/alspac). Briefly, 14,541 pregnant women residents in the former region of Avon, UK, with an expected delivery date between 1^st^ April 1991 and 31^st^ December 1992, were eligible to take part in ALSPAC. Detailed information and biosamples have been collected on these women and their offspring at regular intervals, which are available through a searchable data dictionary (http://www.bris.ac.uk/alspac/researchers/data-access/data-dictionary/).

Written informed consent was obtained for all study participants. Ethical approval for the study was obtained from the ALSPAC Ethics and Law Committee and the Local Research Ethics Committees.

### Accessible Resource for Integrated Epigenomic Studies project (ARIES)

#### Samples

Blood samples were obtained for 1,018 mother-offspring pairs (mothers at two timepoints and their offspring at three timepoints) as part of the Accessible Resource for Integrated Epigenomic Studies project (ARIES)(Relton et al., 2015). The Illumina HumanMethylation450 (450K) BeadChip array was used to measure DNA methylation at over 480,000 sites across the epigenome.

#### Methylation assays

DNA samples were bisulfite treated using the Zymo EZ DNA Methylation^TM^ kit (Zymo, Irvine, CA). The Illumina HumanMethylation450 BeadChip (HM450k) was used to measure methylation across the genome and the following arrays were scanned using Illumina iScan, along with an initial quality review using GenomeStudio. A purpose-built laboratory information management system (LIMS) was responsible for generating batch variables during data generation. LIMS also reported quality control (QC) metrics for the standard probes on the HM450k for all samples and excluded those which failed QC. Data points with a read count of 0 or with low signal:noise ratio (based on a p-value > 0.01) were also excluded based on the QC report from Illumina to maintain the integrity of probe measurements. Methylation measurements were then compared across timepoints for the same individual and with SNP-chip data (HM450k probes clustered using k-means) to identify and remove sample mismatches. All remaining data from probes was normalised with the Touleimat and Tost(Touleimat and Tost, 2012) algorithms using R with the 15atermelon package(Pidsley et al., 2013). This was followed by rank-normalising the data to remove outliers. Potential batch effect were removed by regressing data points on all covariates. These included the bisulfite-converted DNA (BCD) plate batch and white blood cell count which was adjusted for using the *estimateCellCounts* function in the minfi, Bioconductor package(Jaffe and Irizarry, 2014).

#### Genotyping assays

Genotype data were available for all ALSPAC individuals enrolled in the ARIES project, which had previously undergone quality control, cleaning and imputation at the cohort level. ALSPAC offspring selected for this project had previously been genotyped using the Illumina HumanHap550 quad genome-wide SNP genotyping platform (Illumina Inc, San Diego, USA) by the Wellcome Trust Sanger Institute (WTSI, Cambridge, UK) and the Laboratory Corporation of America (LCA, Burlington, NC, USA). Samples were excluded based on incorrect sex assignment; abnormal heterozygosity (<0.320 or >0.345 for WTSI data; <0.310 or >0.330 for LCA data); high missingness (>3%); cryptic relatedness (>10% identity by descent) and non-European ancestry (detected by multidimensional scaling analysis). After QC, 500,527 SNP loci were available for the directly genotyped dataset. Following QC the final directly genotyped dataset contained 526,688 SNP loci.

#### Imputation

Genotypes with MAF > 0.01 and Hardy-Weinberg equilibrium P > 5×10^−7^ were phased together using ShapeIt (version 2, revision 727)(Delaneau et al., 2013) and imputed using the 1000 Genomes reference panel (phase 1, version 3, phased using ShapeIt version 2, December 2013, using all populations) using Impute (v2.2.2)(Howie et al., 2009). After imputation dosages were converted to bestguess genotypes and filtered to only keep variants with an imputation quality score ≥ 0.8. The final imputed dataset used for the analyses presented here contained 8,074,398 loci.

### The mQTL database

Observed effects for genetic variants strongly associated with DNA methylation (referred to hereafter as mQTL) were obtained from the mQTL database (http://www.mqtldb.org/) (Gaunt et al., 2016). In this study we have only used mQTL acting in cis (i.e. variants located within 1MB of their associated CpG site) to reduce the risk of pleiotropy influencing our results, as variants which are associated with methylation levels at multiple loci across the genome may be more likely to impact independent biological pathways simultaneously.

LD clumping was undertaken to identify independent mQTL for each CpG site which could be used as instrumental variables for Mendelian randomization (MR) analyses. In total, there were 30,328 CpG sites eligible for analysis (26,975 CpG sites with 1 mQTL, 5,984 CpG sites with 2 mQTLs, 969 CpG sites with 3 mQTLs, 140 CpG sites with 4 mQTLs and 3 CpG sites with 5 mQTLs). If an mQTL and associated CpG site were observed at more than one of the 5 possible time points measured in the same individuals within ARIES, we used effect estimates from the time point with the largest effect based on p-values.

### GWAS summary data for 139 complex traits and diseases

We identified observed effects for genetic variants on complex traits using large-scale studies which were available within the MR-Base platform (www.mrbase.org) (Hemani et al., 2016). We used the following inclusion criteria to select complex traits to be analysed:

- Effects reported genome-wide for over 95,000 genetic variants
- Study samples must be larger than 1000
- Either European or mixed populations
- Reported beta, standard error and effect alleles for variants

### The UK Biobank

Genotype data was available for approximately 490,000 individuals enrolled in the UK Biobank study. Phasing and imputation of this data is explained elsewhere (Bycroft et al., 2017). Individuals with withdrawn consent, evidence of genetic relatedness or who were not of ‘white European ancestry’ based on a K-means clustering (K=4) were excluded from analysis.

Phenotype data were collected for the following traits (with their UK Biobank variable ID in brackets) which were identified as suitable for replication due to their samples sizes after merging with genotype data (n > 1000); Age at menarche (2714), Age at menopause (3581), Asthma (22127), Birth weight (20022), Body mass index (21001), Cigarettes smoked per day (3456), Extreme Height (derived from 50), Height (50), Hip circumference (49), Myocardial infarction (41202, ICD10 code = I21 or I22), Obesity class 1 (derived from 21001), Type 2 Diabetes (derived from 2443, although this variable does not distinguish between diabetes type), Waist circumference (48), Weight (21002) and Years of schooling (derived from 6138 to calculate EduYears as described by Okbay et al (Okbay et al., 2016)). After exclusions there were up to 334,398 individuals with both genotype and phenotype data who were eligible for analysis.

### Statistical Analysis

#### Identifying candidate loci for mediation by DNA methylation

2SMR was undertaken systematically to evaluate evidence of a causal relationship between DNA methylation at all eligible CpG sites and complex traits. In this initial analysis DNA methylation was treated as our exposure and complex traits as our outcome, using mQTL as our instrumental variables. We used the PhenoSpD method (Zheng et al., 2017, Nyholt, 2004, Cichonska et al., 2016) to calculate the appropriate number of independent traits to adjust our analysis for due to strong correlation amongst certain traits (i.e. BMI and obesity). The multiple testing threshold was calculated as 0.05 divided by the derived number of independent tests. CpG sites for effects which survived this threshold were annotated based on evaluations of the 450K array (Naeem et al., 2014, Zhou et al., 2017). When only one valid genetic instrument was available MR effect estimates are based on the Wald ratio test. Where two or more valid genetic instruments were available for analysis we used the inverse variance weighted (IVW) method to obtain MR effect estimates(Lawlor et al., 2008). Results were plotted as Manhattan plots using code derived from the qqman package in R (Turner, 2014).

#### Distinguishing causal effects from genetic confounding due to linkage disequilibrium

Results which survived the multiple testing threshold in the previous analyses were evaluated using the joint likelihood method (JLIM) (Chun et al., 2017). The JLIM method evaluates whether the same underlying genetic variation is responsible for observed effects on two traits (i.e. DNA methylation at a CpG site and a complex trait in this study). This is achieved using individual-level data for one trait, which was DNA methylation levels obtained from the ARIES project in this study, to generate a permutation-based null distribution. The number of permutations required by the JLIM method was determined by number of tests undertaken (i.e. the number of effects which survived the p-value threshold in the previous analysis). A lack of evidence (i.e. P < 0.05/number of effects evaluated) in this analysis would suggest that the causal variant for methylation variation was simply in linkage disequilibrium with the putative causal variant for the trait (thus introducing genetic confounding into the association between DNA methylation and complex trait).

The JLIM approach was selected over alternative co-localization methods (such as the HEIDI (heterogeneity in dependent instruments)(Zhu et al., 2016) and ‘coloc’ methods(Giambartolomei et al., 2014)) as in this study we always had individual-level data for one of the traits being assessed (epigenome-wide DNA methylation levels from the ARIES project) and therefore did not have to rely on availability of summary statistics for both traits. The authors of the JLIM method also demonstrate strong overall performance compared to alternative approaches, although they do specify two limitations to ensure accurate detection of shared genetic effects between two traits. These limitations are that their resolution becomes limited when 1) at high LD levels (i.e. r^2^ ≥ 0.8) between multiple causal instruments and 2) when the QTL effect (i.e. mQTL in this study) is very weak (i.e. P > 0.01). These were addressed in our study as we only used multiple instruments within the MR analysis that were independent (r^2^ < 0.01) and strongly associated with DNA methylation (P < 1.0 × 10^−7^).

#### Reverse Mendelian randomization

For CpG-trait effects identified in the previous analysis, we also used 2SMR to evaluate evidence of genetic liability by modelling complex traits as our exposure and DNA methylation as our outcome. Instruments for complex traits were selected based on a threshold of 5.0 × 10^−08^ from large-scale GWAS after LD clumping to identify independent variants. The IVW method was applied to estimate the causal effects of traits on CpG sites where more than one instrument was available, otherwise the Wald ratio was used.

#### Replication of observed effects in UK Biobank

For CpG-trait effects where DNA methylation and complex trait were driven by the same causal variant, as inferred by the JLIM method, we repeated our initial analysis using data from the UK Biobank project(Sudlow et al., 2015). Therefore, our estimates of genetic variants on complex trait variation have been obtained in a separate population in these analyses, whereas estimates on DNA methylation remain the same as in the discovery analysis as there is currently no appropriate replication sample.

This validation analysis was undertaken for effects across 14 traits from the full release of the UK Biobank project for which large sample sizes (n ≥ 10,000) were available after merging with available genetic data (Table S4) (Sudlow et al., 2015). Linear or logistic regression was used (depending on whether the trait was continuous or binary respectively) to determine effect estimates of genetic variants on complex traits adjusted for age, sex, the first 10 principal components and a binary indicator which reflects which genotype chip individuals were measured on. This was because a subset of UK Biobank individuals had their genotype measured on the Affymetrix UK BiLEVE Axiom array (~50,000 participants), whereas the remainder were measured using the Affymetrix UK Biobank Axiom array.

#### Causal relationship between DNA methylation and gene expression

We undertook 2SMR to evaluate the relationship between DNA methylation and gene expression for effects where the causal variant, as indicated by the JLIM method described above, was both an mQTL and eQTL. Effect estimates for variants on gene expression were obtained from the GTEx consortium (http://www.gtexportal.org/)(Consortium, 2013). When effect estimates for the putative causal variant were not available from GTEx we identified a surrogate variant instead (r^2^≥0.8). Where no surrogate was available within GTEx we consulted the blood eQTL browser (http://genenetwork.nl/bloodeqtlbrowser/)(Westra et al., 2013).

### Functional informatics

#### Variant annotation and gene prioritisation

Genetic variants for effects potentially mediated by changes in DNA methylation were analysed using the variant effect predictor (VEP)(McLaren et al., 2016) to calculate their predicted consequence. Regulatory data were obtained from Ensembl (www.ensembl.org/)(Yates et al., 2016) to evaluate whether these variants reside within regulatory regions of the genome.

Prior to enrichment analyses and gene prioritization, as effects were grouped together as opposed to evaluated individually, we removed observed effects involving CpG sites flagged for exclusion based on evaluations by Naeem et al (Naeem et al., 2014). This was based on their criteria of overlapping SNPs at CpG probes, probes which map to multiple locations and repeats on the 450K array. The DEPICT method (Pers et al., 2015) was used to prioritise genes for all remaining variants. Variants which were not allocated a likely impacted gene by DEPICT were annotated with their nearest gene using bedtools (Quinlan, 2014).

#### Pathway and gene ontology enrichment

Genes implicated in the previous evaluations were tested for enrichment of functional pathways and gene ontology terms using ConsensusPathDB (Kamburov et al., 2013). When multiple genes were implicated at the same association signal we used annotations according to DEPICT over the nearest gene. All results which had a false discovery rate < 5% were reported.

#### Identifying known and candidate genes for therapeutic intervention

We consulted the ChEMBL database (Bento et al., 2014) (version 23 accessed on 13^th^ June 2017) to ascertain whether any of the implicated genes encode proteins for known targets of approved drugs or compounds in development.

#### Tissue specific enrichment for CpG sites

The hypergeometric test was used to test for enrichment of implicated CpG sites for histone mark peaks and regions of DNAse I in up to 113 different tissue and cell types from the Encyclopedia of DNA Elements (ENCODE) and Roadmap Epigenomics projects. To calibrate background expectations, we randomly selected CpG sites across the epigenome which resided in similar genomic regions based on Illumina annotations (i.e. CpG island, gene body etc.). We used permutations to control for multiple testing by randomly selecting the same number of implicated CpG sites matched on location and then repeating the enrichment computation for 10,000 iterations. This analysis was repeated using regulatory annotations from the Illumina 450K file (enhancer regions) and Ensembl (promoters, open chromatin regions, transcriptional repressor CTCF sites and transcription factor binding sites).

## Acknowledgements

We are extremely grateful to all the families who took part in this study, the midwives for their help in recruiting them, and the whole ALSPAC team, which includes interviewers, computer and laboratory technicians, clerical workers, research scientists, volunteers, managers, receptionists and nurses. The UK Medical Research Council and the Wellcome Trust (Grant ref: 102215/2/13/2) and the University of Bristol provide core support for ALSPAC. GWAS data was generated by Sample Logistics and Genotyping Facilities at the Wellcome Trust Sanger Institute and LabCorp (Laboratory Corporation of America) using support from 23andMe. Methylation data in the ALSPAC cohort was generated as part of the UK BBSRC funded (BB/I025751/1 and BB/I025263/1) Accessible Resource for Integrated Epigenomic Studies (ARIES). UK Biobank data was analysed as part of projects 8786 and 15825.

This publication is the work of the authors and Tom G Richardson will serve as guarantor for the contents of this paper. This work was supported by the UK Medical Research Council (MRC Integrative Epidemiology Unit, MC UU 12013/1, MC UU 12013/2, MC UU 12013/3, MC UU 12013/8). NJT is a Wellcome Trust Investigator (202802/Z/16/Z), is a programme lead in the MRC Integrative Epidemiology Unit (MC_UU_12013/3) and works within the University of Bristol NIHR Biomedical Research Centre (BRC). TGR is supported by the Elizabeth Blackwell Institute Proximity to Discovery award (EBI 424). The authors declare no conflicts of interest.

## References

Bento, A. P., Gaulton, A., Hersey, A., Bellis, L. J., Chambers, J., Davies, M., Kruger, F. A., Light, Y., Mak, L., Mcglinchey, S., Nowotka, M., Papadatos, G., Santos, R. & Overington, J. P. 2014. The ChEMBL bioactivity database: an update. Nucleic Acids Res, 42, D1083–90.

Bowden, J., Davey Smith, G. & Burgess, S. 2015. Mendelian randomization with invalid instruments: effect estimation and bias detection through Egger regression. Int J Epidemiol, 44, 512–25.

Bowden, J., Davey Smith, G., Haycock, P. C. & Burgess, S. 2016. Consistent Estimation in Mendelian Randomization with Some Invalid Instruments Using a Weighted Median Estimator. Genet Epidemiol, 40, 304–14.

Boyd, A., Golding, J., Macleod, J., Lawlor, D. A., Fraser, A., Henderson, J., Molloy, L., Ness, A., Ring, S. & Davey Smith, G. 2013. Cohort Profile: the ‘children of the 90s’–the index offspring of the Avon Longitudinal Study of Parents and Children. Int J Epidemiol, 42, 111–27.

Burgess, S., Scott, R. A., Timpson, N. J., Davey Smith, G., Thompson, S. G. & Consortium, E.-I. 2015. Using published data in Mendelian randomization: a blueprint for efficient identification of causal risk factors. Eur J Epidemiol, 30, 543–52.

Burkhardt, R., Kirsten, H., Beutner, F., Holdt, L. M., Gross, A., Teren, A., Tonjes, A., Becker, S., Krohn, K., Kovacs, P., Stumvoll, M., Teupser, D., Thiery, J., Ceglarek, U. & Scholz, M. 2015. Integration of Genome-Wide SNP Data and Gene-Expression Profiles Reveals Six Novel Loci and Regulatory Mechanisms for Amino Acids and Acylcarnitines in Whole Blood. PLoS Genet, 11, e1005510.

Bycroft, C., Freeman, C., Petkova, D., Band, G., Elliott, L. T., Sharp, K., Motyer, A., Vukcevic, D., Delaneau, G., O’Connell, J., Cortes, A., Welsh, S., Mcvean, G., Leslie, S., Donnelly, P. & Marchini, J. 2017. Genome-wide genetic data on ~500,000 UK Biobank participants. http://www.biorxiv.org/content/early/2017/07/20/166298.

Carithers, L. J. & Moore, H. M. 2015. The Genotype-Tissue Expression (GTEx) Project. Biopreserv Biobank, 13, 307–8.

Chun, S., Casparino, A., Patsopoulos, N. A., Croteau-Chonka, D. C., Raby, B. A., De Jager, P. L., Sunyaev, S. R. & Cotsapas, C. 2017. Limited statistical evidence for shared genetic effects of eQTLs and autoimmune-disease-associated loci in three major immune-cell types. Nat Genet, 49, 600–605.

Cichonska, A., Rousu, J., Marttinen, P., Kangas, A. J., Soininen, P., Lehtimaki, T., Raitakari, O. T., Jarvelin, M. R., Salomaa, V., Ala-Korpela, M., Ripatti, S. & Pirinen, M. 2016. metaCCA: summary statistics-based multivariate meta-analysis of genome-wide association studies using canonical correlation analysis. Bioinformatics, 32, 1981–9.

Consortium, G. T. 2013. The Genotype-Tissue Expression (GTEx) project. Nat Genet, 45, 580–5.

Davey Smith, G. & Ebrahim, S. 2003. ’Mendelian randomization’: can genetic epidemiology contribute to understanding environmental determinants of disease? Int J Epidemiol, 32, 1–22.

Davey Smith, G. & Hemani, G. 2014. Mendelian randomization: genetic anchors for causal inference in epidemiological studies. Hum Mol Genet, 23, R89–98.

Delaneau, O., Howie, B., Cox, A. J., Zagury, J. F. & Marchini, J. 2013. Haplotype estimation using sequencing reads. Am J Hum Genet, 93, 687–96.

Edwards, S. L., Beesley, J., French, J. D. & Dunning, A. M. 2013. Beyond GWASs: illuminating the dark road from association to function. Am J Hum Genet, 93, 779–97.

Encode Project Consortium, Birney, E., Stamatoyannopoulos, J. A., Dutta, A., Guigo, R., Gingeras, T. R., Margulies, E. H., Weng, Z., Snyder, M., Dermitzakis, E. T., Thurman, R. E., Kuehn, M. S., Taylor, C. M., Neph, S., Koch, C. M., Asthana, S., Malhotra, A., Adzhubei, I., Greenbaum, J. A., Andrews, R. M., Flicek, P., Boyle, P. J., Cao, H., Carter, N. P., Clelland, G. K., Davis, S., Day, N., Dhami, P., Dillon, S. C., Dorschner, M. O., Fiegler, H., Giresi, P. G., Goldy, J., Hawrylycz, M., Haydock, A., Humbert, R., James, K. D., Johnson, B. E., Johnson, E. M., Frum, T. T., Rosenzweig, E. R., Karnani, N., Lee, K., Lefebvre, G. C., Navas, P. A., Neri, F., Parker, S. C., Sabo, P. J., Sandstrom, R., Shafer, A., Vetrie, D., Weaver, M., Wilcox, S., Yu, M., Collins, F. S., Dekker, J., Lieb, J. D., Tullius, T. D., Crawford, G. E., Sunyaev, S., Noble, W. S., Dunham, I., Denoeud, F., Reymond, A., Kapranov, P., Rozowsky, J., Zheng, D., Castelo, R., Frankish, A., Harrow, J., Ghosh, S., Sandelin, A., Hofacker, I. L., Baertsch, R., Keefe, D., Dike, S., Cheng, J., Hirsch, H. A., Sekinger, E. A., Lagarde, J., Abril, J. F., Shahab, A., Flamm, C., Fried, C., Hackermuller, J., Hertel, J., Lindemeyer, M., Missal, K., Tanzer, A., Washietl, S., Korbel, J., Emanuelsson, O., Pedersen, J. S., Holroyd, N., Taylor, R., Swarbreck, D., Matthews, N., Dickson, M. C., Thomas, D. J., Weirauch, M. T., et al. 2007. Identification and analysis of functional elements in 1% of the human genome by the ENCODE pilot project. Nature, 447, 799–816.

Fraser, A., Macdonald-Wallis, C., Tilling, K., Boyd, A., Golding, J., Davey Smith, G., Henderson, J., Macleod, J., Molloy, L., Ness, A., Ring, S., Nelson, S. M. & Lawlor, D. A. 2013. Cohort Profile: the Avon Longitudinal Study of Parents and Children: ALSPAC mothers cohort. Int J Epidemiol, 42, 97–110.

Gaunt, T. R., Shihab, H. A., Hemani, G., Min, J. L., Woodward, G., Lyttleton, O., Zheng, J., Duggirala, A., Mcardle, W. L., Ho, K., Ring, S. M., Evans, D. M., Davey Smith, G. & Relton, C. L. 2016. Systematic identification of genetic influences on methylation across the human life course. Genome Biol, 17, 61.

Giambartolomei, C., Vukcevic, D., Schadt, E. E., Franke, L., Hingorani, A. D., Wallace, C. & Plagnol, V. 2014. Bayesian test for colocalisation between pairs of genetic association studies using summary statistics. PLoS Genet, 10, e1004383.

Hannon, E., Weedon, M., Bray, N., O’Donovan, M. & Mill, J. 2017. Pleiotropic Effects Of Trait-Associated Genetic Variation on DNA Methylation: Utility for Refining GWAS Loci. Am J Hum Genet, 100, 954–959.

Hartwig, F. P., Davey Smith, G. & Bowden, J. 2017. Robust Inference In Two-Sample Mendelian Randomisation Via The Zero Modal Pleiotropy Assumption. http://www.biorxiv.org/content/early/2017/04/10/126102.

Hemani, G., Zheng, J., Wade, K. H., Laurin, C., Elsworth, E., Burgess, S., Bowden, J., Langdon, R., Tan, V., Yarmolinsky, J., Shihab, H. A., Timpson, N., Evans, D. M., Relton, C. L., Martin, R., Davey Smith, G., Gaunt, T. & Haycock, P. C. 2016. MR-Base: a platform for systematic causal inference across the phenome using billions of genetic associations.

Howie, B. N., Donnelly, P. & Marchini, J. 2009. A Flexible And Accurate Genotype Imputation Method For the next generation of genome-wide association studies. PLoS Genet, 5, e1000529.

Jaffe, A. E. & Irizarry, R. A. 2014. Accounting for cellular heterogeneity is critical in epigenome-wide association studies. Genome Biol, 15, R31.

Joehanes, R., Just, A. C., Marioni, R. E., Pilling, L. C., Reynolds, L. M., Mandaviya, P. R., Guan, W., Xu, T., Elks, C. E., Aslibekyan, S., Moreno-Macias, H., Smith, J. A., Brody, J. A., Dhingra, R., Yousefi, P., Pankow, J. S., Kunze, S., Shah, S. H., Mcrae, A. F., Lohman, K., Sha, J., Absher, D. M., Ferrucci, L., Zhao, W., Demerath, E. W., Bressler, J., Grove, M. L., Huan, T., Liu, C., Mendelson, M. M., Yao, C., Kiel, D. P., Peters, A., Wang-Sattler, R., Visscher, P. M., Wray, N. R., Starr, J. M., Ding, J., Rodriguez, C. J., Wareham, N. J., Irvin, M. R., Zhi, D., Barrdahl, M., Vineis, P., Ambatipudi, S., Uitterlinden, A. G., Hofman, A., Schwartz, J., Colicino, E., Hou, L., Vokonas, P. S., Hernandez, D. G., Singleton, A. B., Bandinelli, S., Turner, S. T., Ware, E. B., Smith, A. K., Klengel, T., Binder, E. B., Psaty, B. M., Taylor, K. D., Gharib, S. A., Swenson, B. R., Liang, L., Demeo, D. L., O’Connor, G. T., Herceg, Z., Ressler, K. J., Conneely, K. N., Sotoodehnia, N., Kardia, S. L., Melzer, D., Baccarelli, A. A., Van Meurs, J. B., Romieu, I., Arnett, D. K., Ong, K. K., Liu, Y., Waldenberger, M., Deary, I. J., Fornage, M., Levy, D. & London, S. J. 2016. Epigenetic Signatures of Cigarette Smoking. Circ Cardiovasc Genet, 9, 436–447.

Joubert, B. R., Felix, J. F., Yousefi, P., Bakulski, K. M., Just, A. C., Breton, C., Reese, S. E., Markunas, C. A., Richmond, R. C., Xu, C. J., Kupers, L. K., Oh, S. S., Hoyo, C., Gruzieva, O., Soderhall, C., Salas, L. A., Baiz, N., Zhang, H., Lepeule, J., Ruiz, C., Ligthart, S., Wang, T., Taylor, J. A., Duijts, L., Sharp, G. C., Jankipersadsing, S. A., Nilsen, R. M., Vaez, A., Fallin, M. D., Hu, D., Litonjua, A. A., Fuemmeler, B. F., Huen, K., Kere, J., Kull, I., Munthe-Kaas, M. C., Gehring, U., Bustamante, M., Saurel-Coubizolles, M. J., Quraishi, B. M., Ren, J., Tost, J., Gonzalez, J. R., Peters, M. J., Haberg, S. E., Xu, Z., Van Meurs, J. B., Gaunt, T. R., Kerkhof, M., Corpeleijn, E., Feinberg, A. P., Eng, C., Baccarelli, A. A., Benjamin Neelon, S. E., Bradman, A., Merid, S. K., Bergstrom, A., Herceg, Z., Hernandez-Vargas, H., Brunekreef, B., Pinart, M., Heude, B., Ewart, S., Yao, J., Lemonnier, N., Franco, O. H., Wu, M. C., Hofman, A., Mcardle, W., Van Der Vlies, P., Falahi, F., Gillman, M. W., Barcellos, L. F., Kumar, A., Wickman, M., Guerra, S., Charles, M. A., Holloway, J., Auffray, C., Tiemeier, H. W., Smith, G. D., Postma, D., Hivert, M. F., Eskenazi, B., Vrijheid, M., Arshad, H., Anto, J. M., Dehghan, A., Karmaus, W., Annesi-Maesano, I., Sunyer, J., Ghantous, A., Pershagen, G., Holland, N., Murphy, S. K., Demeo, D. L., Burchard, E. G., Ladd-Acosta, C., Snieder, H., Nystad, W., et al. 2016. DNA Methylation in Newborns and Maternal Smoking in Pregnancy: Genome-wide Consortium Meta-analysis. Am J Hum Genet, 98, 680–96.

Kamburov, A., Stelzl, U., Lehrach, H. & Herwig, R. 2013. The ConsensusPathDB interaction database: 2013 update. Nucleic Acids Res, 41, D793–800.

Lawlor, D. A., Harbord, R. M., Sterne, J. A., Timpson, N. & Davey Smith, G. 2008. Mendelian randomization: using genes as instruments for making causal inferences in epidemiology. Stat Med, 27, 1133–63.

Linnér, R. K. et al. 2017. An epigenome-wide association study of educational attainment (n = 10,767). http://biorxiv.org/content/early/2017/03/07/114637.

Ma, X., Wang, Y. W., Zhang, M. Q. & Gazdar, A. F. 2013. DNA methylation data analysis and its application to cancer research. Epigenomics, 5, 301–16.

Mancuso, N., Shi, H., Goddard, P., Kichaev, G., Gusev, A. & Pasaniuc, B. 2017. Integrating Gene Expression with Summary Association Statistics to Identify Genes Associated with 30 Complex Traits. Am J Hum Genet, 100, 473–487.

Mclaren, W., Gil, L., Hunt, S. E., Riat, H. S., Ritchie, G. R., Thormann, A., Flicek, P. & Cunningham, F. 2016. The Ensembl Variant Effect Predictor. Genome Biol, 17, 122.

Mcrae, A. F., Powell, J. E., Henders, A. K., Bowdler, L., Hemani, G., Shah, S., Painter, J. N., Martin, N. G., Visscher, P. M. & Montgomery, G. W. 2014. Contribution of genetic variation to transgenerational inheritance of DNA methylation. Genome Biol, 15, R73.

Naeem, H., Wong, N. C., Chatterton, Z., Hong, M. K., Pedersen, J. S., Corcoran, N. M., Hovens, C. M. & Macintyre, G. 2014. Reducing the risk of false discovery enabling identification of biologically significant genome-wide methylation status using the HumanMethylation450 array. BMC Genomics, 15, 51.

Nyholt, D. R. 2004. A simple correction for multiple testing for single-nucleotide polymorphisms in linkage disequilibrium with each other. Am J Hum Genet, 74, 765–9.

Okbay, A., Beauchamp, J. P., Fontana, M. A., Lee, J. J., Pers, T. H., Rietveld, C. A., Turley, P., Chen, G. B., Emilsson, V., Meddens, S. F., Oskarsson, S., Pickrell, J. K., Thom, K., Timshel, P., De Vlaming, R., Abdellaoui, A., Ahluwalia, T. S., Bacelis, J., Baumbach, C., Bjornsdottir, G., Brandsma, J. H., Pina Concas, M., Derringer, J., Furlotte, N. A., Galesloot, T. E., Girotto, G., Gupta, R., Hall, L. M., Harris, S. E., Hofer, E., Horikoshi, M., Huffman, J. E., Kaasik, K., Kalafati, I. P., Karlsson, R., Kong, A., Lahti, J., Van Der Lee, S. J., Deleeuw, C., Lind, P. A., Lindgren, K. O., Liu, T., Mangino, M., Marten, J., Mihailov, E., Miller, M. B., Van Der Most, P. J., Oldmeadow, C., Payton, A., Pervjakova, N., Peyrot, W. J., Qian, Y., Raitakari, O., Rueedi, R., Salvi, E., Schmidt, B., Schraut, K. E., Shi, J., Smith, A. V., Poot, R. A., St Pourcain, B., Teumer, A., Thorleifsson, G., Verweij, N., Vuckovic, D., Wellmann, J., Westra, H. J., Yang, J., Zhao, W., Zhu, Z., Alizadeh, B. Z., Amin, N., Bakshi, A., Baumeister, S. E., Biino, G., Bonnelykke, K., Boyle, P. A., Campbell, H., Cappuccio, F. P., Davies, G., De Neve, J. E., Deloukas, P., Demuth, I., Ding, J., Eibich, P., Eisele, L., Eklund, N., Evans, D. M., Faul, J. D., Feitosa, M. F., Forstner, A. J., Gandin, I., Gunnarsson, B., Halldorsson, B. V., Harris, T. B., Heath, A. C., Hocking, L. J., Holliday, E. G., Homuth, G., Horan, M. A., et al. 2016. Genome-wide association study identifies 74 loci associated with educational attainment. Nature, 533, 539–42.

Pers, T. H., Karjalainen, J. M., Chan, Y., Westra, H. J., Wood, A. R., Yang, J., Lui, J. C., Vedantam, S., Gustafsson, S., Esko, T., Frayling, T., Speliotes, E. K., Genetic Investigation Of, A. T.C., Boehnke, M., Raychaudhuri, S., Fehrmann, R. S., Hirschhorn, J. N. & Franke, L. 2015. Biological interpretation of genome-wide association studies using predicted gene functions. Nat Commun, 6, 5890.

Pidsley, R., Y Wong, C. C., Volta, M., Lunnon, K., Mill, J. & Schalkwyk, L. C. 2013. A data-driven approach to preprocessing Illumina 450K methylation array data. BMC Genomics, 14, 293.

Pierce, B. L. & Burgess, S. 2013. Efficient design for Mendelian randomization studies: subsample and 2-sample instrumental variable estimators. Am J Epidemiol, 178, 1177–84.

Quinlan, A. R. 2014. BEDTools: The Swiss-Army Tool for Genome Feature Analysis. Curr Protoc Bioinformatics, 47, 11 12 1–34.

Relton, C. L. & Davey Smith, G. 2010. Epigenetic epidemiology of common complex disease: prospects for prediction, prevention, and treatment. PLoS Med, 7, e1000356.

Relton, C. L., Gaunt, T., Mcardle, W., Ho, K., Duggirala, A., Shihab, H., Woodward, G., Lyttleton, O., Evans, D. M., Reik, W., Paul, Y. L., Ficz, G., Ozanne, S. E., Wipat, A., Flanagan, K., Lister, A., Heijmans, B. T., Ring, S. M. & Davey Smith, G. 2015. Data Resource Profile: Accessible Resource for Integrated Epigenomic Studies (ARIES). Int J Epidemiol.

Richardson, T. G., Zheng, J., Davey Smith, G., Timpson, N. J., Gaunt, T. R., Relton, C. L. & Hemani, G. 2017. Causal epigenome-wide association study identifies CpG sites that influence cardiovascular disease risk. http://biorxiv.org/content/early/2017/04/29/132019 (due to appear in the American Journal of Human Genetics).

Romanoski, C. E., Glass, C. K., Stunnenberg, H. G., Wilson, L. & Almouzni, G. 2015. Epigenomics: Roadmap for regulation. Nature, 518, 314–6.

Sudlow, C., Gallacher, J., Allen, N., Beral, V., Burton, P., Danesh, J., Downey, P., Elliott, P., Green, J., Landray, M., Liu, B., Matthews, P., Ong, G., Pell, J., Silman, A., Young, A., Sprosen, T., Peakman, T. & Collins, R. 2015. UK biobank: an open access resource for identifying the causes of a wide range of complex diseases of middle and old age. PLoS Med, 12, e1001779.

Timpson, N. J., Nordestgaard, B. G., Harbord, R. M., Zacho, J., Frayling, T. M., Tybjaerg-Hansen, A. & Smith, G. D. 2011. C-reactive protein levels and body mass index: elucidating direction of causation through reciprocal Mendelian randomization. Int J Obes (Lond), 35, 300–8.

Touleimat, N. & Tost, J. 2012. Complete pipeline for Infinium(®) Human Methylation 450K BeadChip data processing using subset quantile normalization for accurate DNA methylation estimation. Epigenomics, 4, 325–41.

Turner, S. D. 2014. qqman: an R package for visualizing GWAS results using Q-Q and manhattan plots.

Vimaleswaran, K. S., Berry, D. J., Lu, C., Tikkanen, E., Pilz, S., Hiraki, L. T., Cooper, J. D., Dastani, Z., Li, R., Houston, D. K., Wood, A. R., Michaelsson, K., Vandenput, L., Zgaga, L., Yerges-Armstrong, L. M., Mccarthy, M. I., Dupuis, J., Kaakinen, M., Kleber, M. E., Jameson, K., Arden, N., Raitakari, O., Viikari, J., Lohman, K. K., Ferrucci, L., Melhus, H., Ingelsson, E., Byberg, L., Lind, L., Lorentzon, M., Salomaa, V., Campbell, H., Dunlop, M., Mitchell, B. D., Herzig, K. H., Pouta, A., Hartikainen, A. L., Genetic Investigation Of Anthropometric Traits, G. C., Streeten, E. A., Theodoratou, E., Jula, A., Wareham, N. J., Ohlsson, C., Frayling, T. M., Kritchevsky, S. B., Spector, T. D., Richards, J. B., Lehtimaki, T., Ouwehand, W. H., Kraft, P., Cooper, C., Marz, W., Power, C., Loos, R. J., Wang, T. J., Jarvelin, M. R., Whittaker, J. C., Hingorani, A. D. & Hypponen, E. 2013. Causal relationship between obesity and vitamin D status: bi-directional Mendelian randomization analysis of multiple cohorts. PLoS Med, 10, e1001383.

Westra, H. J., Peters, M. J., Esko, T., Yaghootkar, H., Schurmann, C., Kettunen, J., Christiansen, M. W., Fairfax, B. P., Schramm, K., Powell, J. E., Zhernakova, A., Zhernakova, D. V., Veldink, J. H., Van Den Berg, L. H., Karjalainen, J., Withoff, S., Uitterlinden, A. G., Hofman, A., Rivadeneira, F., T Hoen, P. A., Reinmaa, E., Fischer, K., Nelis, M., Milani, L., Melzer, D., Ferrucci, L., Singleton, A. B., Hernandez, D. G., Nalls, M. A., Homuth, G., Nauck, M., Radke, D., Volker, U., Perola, M., Salomaa, V., Brody, J., Suchy-Dicey, A., Gharib, S. A., Enquobahrie, D. A., Lumley, T., Montgomery, G. W., Makino, S., Prokisch, H., Herder, C., Roden, M., Grallert, H., Meitinger, T., Strauch, K., Li, Y., Jansen, R. C., Visscher, P. M., Knight, J. C., Psaty, B. M., Ripatti, S., Teumer, A., Frayling, T. M., Metspalu, A., Van Meurs, J. B. & Franke, L. 2013. Systematic identification of trans eQTLs as putative drivers of known disease associations. Nat Genet, 45, 1238–43.

Yates, A., Akanni, W., Amode, M. R., Barrell, D., Billis, K., Carvalho-Silva, D., Cummins, C., Clapham, P., Fitzgerald, S., Gil, L., Giron, C. G., Gordon, L., Hourlier, T., Hunt, S. E., Janacek, S. H., Johnson, N., Juettemann, T., Keenan, S., Lavidas, I., Martin, F. J., Maurel, T., Mclaren, W., Murphy, D. N., Nag, R., Nuhn, M., Parker, A., Patricio, M., Pignatelli, M., Rahtz, M., Riat, H. S., Sheppard, D., Taylor, K., Thormann, A., Vullo, A., Wilder, S. P., Zadissa, A., Birney, E., Harrow, J., Muffato, M., Perry, E., Ruffier, M., Spudich, G., Trevanion, S. J., Cunningham, F., Aken, B. L., Zerbino, D. R. & Flicek, P. 2016. Ensembl 2016. Nucleic Acids Res, 44, D710–6.

Zheng, J., Richardson, T. G., Millard, L., Hemani, G., Raistrick, C., Vilhjalmsson, B., Haycock, P. C. & Gaunt, T. R. 2017. PhenoSpD: an integrated toolkit for phenotypic correlation estimation and multiple testing correction using GWAS summary statistics. http://biorxiv.org/content/early/2017/07/25/148627.

Zhou, W., Laird, P. W. & Shen, H. 2017. Comprehensive characterization, annotation and innovative use of Infinium DNA methylation BeadChip probes. Nucleic Acids Res, 45, e22.

Zhu, Z., Zhang, F., Hu, H., Bakshi, A., Robinson, M. R., Powell, J. E., Montgomery, G. W., Goddard, M. E., Wray, N. R., Visscher, P. M. & Yang, J. 2016. Integration of summary data from GWAS and eQTL studies predicts complex trait gene targets. Nat Genet, 48, 481–7.

